# Grammatical category and the neural processing of phrases

**DOI:** 10.1101/2020.12.22.423994

**Authors:** Amelia Burroughs, Nina Kazanina, Conor Houghton

## Abstract

The interlocking roles of lexical, syntactic and semantic processing in language comprehension has been the subject of longstanding debate. Recently, the cortical response to a frequency-tagged linguistic stimulus has been shown to track the rate of phrase and sentence, as well as syllable, presentation. This could be interpreted as evidence for the hierarchical processing of speech, or as a response to the repetition of grammatical category. To examine the extent to which hierarchical structure plays a role in language processing we recorded EEG from human participants as they listen to isochronous streams of monosyllabic words. Comparing responses to sequences in which grammatical category is strictly alternating and chosen such that two-word phrases can be grammatically constructed — `cold food loud room` — or is absent — `rough give ill tell` — showed cortical entrainment at the two-word phrase rate was only present in the grammatical condition. Thus, grammatical category repetition alone does not yield entertainment at higher level than a word. On the other hand, cortical entrainment was reduced for the mixed-phrase condition that contained two-word phrases but no grammatical category repetition — `that word send less` — which is not what would be expected if the measured entrainment reflected purely abstract hierarchical syntactic units. Our results support a model in which word-level grammatical category information is required to build larger units.

## Introduction

The ability of the human brain to rapidly generate meaning from an incoming stream of words is an impressive feat. The role played by hierarchical syntactic structure during this processing is the subject of an ongoing debate with, on the two extremes, some arguing that full hierarchical analysis is central to sentence comprehension [2, 5, 9], while others claim that hierarchical representations are non-essential [10–13].

According to the hierarchical account of language, comprehension is underpinned by the brain’s ability to abstract over a number of linguistic levels, such as grammatical categories and phrases and to combine them hierarchically according to a set of grammatical principles. In this view language users parse an incoming sequence of words into a nested tree-like structure that details taxonomy-like relationships between syntactic constituents and enables sentence comprehension (Fig. 1).

**Fig 1.**
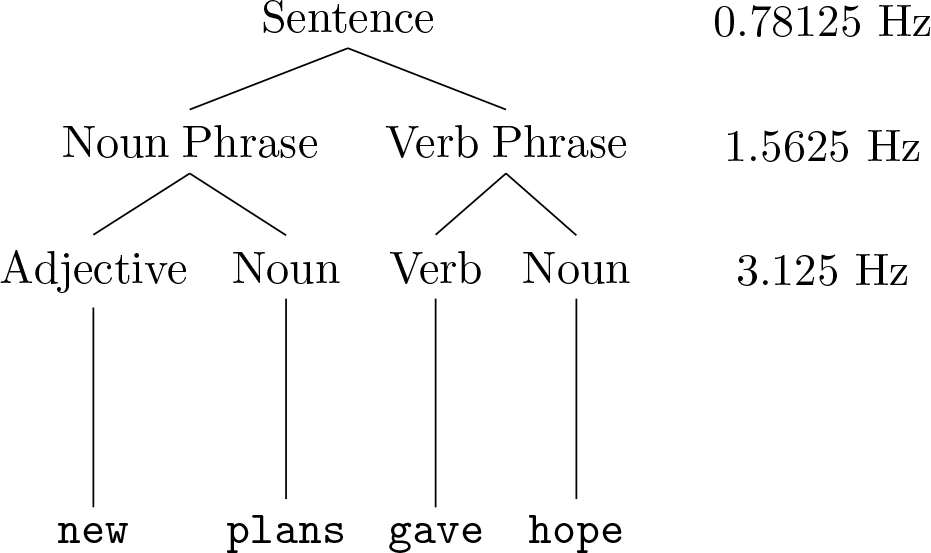
Demonstration of a syntactic tree for an example sentence from [7, 8]. The sentence is composed of a noun phrase and a verb phrase, each consisting of two words presented at a rate of 3.125 Hz. The sentence is described using a hierarchical tree that splits the sentence first into a noun phrase and a verb phrase and then into words. This is a particularly simple tree, more complex trees may have more branches. This simple structure is convenient for frequency tagging and the different frequencies corresponding to the three levels of the tree, sentence, phrase and word, have been marked here.

In support of this view, it has been demonstrated, using MEG in [8] and using EEG in [7] that cortical activity can entrain to the rate of syllable, phrase and sentence presentation. In the EEG experiments participants were played continuous streams of four-word sentences, where each word was 320 ms long in duration and consisted of only a single syllable. As in Fig. 1 each sentence was composed of a noun phrase and a verb phrase, each containing two words. Thus these stimuli have a specific frequency at three levels of linguistic structure: syllables at 3.125 Hz, phrases at 1.5625 Hz and sentences at 0.78125 Hz. The neural responses were analysed using time-frequency decomposition and measures of inter-trial phase coherence (ITPC). Cortical activity was found to be phase-locked to the rate of presentation of syllables, phrases and sentences even though only the syllable frequency was present in the auditory signal itself, the other two frequencies rely on the meaning of the words and the structure of the sentences.

However, it has also been suggested that the brain could rely on simpler, potentially more generic, strategies underpinned by statistical processing of linguistic representations. In line with this, recent work [13] has shown that a model solely based on distributional word semantics is sufficient to predict the response observed in [7, 8]. In this model the distributional word semantics are represented by skipgram-word2vec vectors [4, 17]. In skipgram, word2vec vectors are calculated by training a simple linear neural network with one hidden layer; the input and output layers both correspond to words and the network is trained on the task of predicting, from a given input word, the unordered list of words that occur in proximity to it in text. The components of the word2vec vector for a given word are the weights feeding forwards from the word to the hidden layer. Words that are likely to occur in a similar context have similar representations in the hidden layer and hence are associated with similar word2vec vectors. To a striking degree these high-dimensional vectors have specific directions that serve, at least locally, to represent specific concepts, so that, for example, the same direction that leads from “big” to “biggest” leads from “small” to “smallest” [16, 18].

In [13] fictive EEG signals representing experimental trials were constructed from the word2vec vectors for each stimulus. In the EEG experiment each word was presented for 320 ms and so, in the ficitive data 320 copies of the vector for each word were lined up side-by-side forming the columns of a matrix so each column represents 1 ms of the stimulus. The rows of this matrix were then treated as a EEG and this fictive signal was analysed in the same way as the real EEG signal is, with measures of the evoked response averaged over rows, much as we average over individual electrodes. This simulated EEG signal demonstrated the same entrainment to words, phrases and sentences as the real signal. Since the high-dimensional word2vec vectors represent single words only and do not explicitly encode information about word sequences, this demonstrated that semantic relationships that can be deduced from a text corpus are sufficient to explain the ITPC peaks seen in the real experiment, without any need to invoke the hierarchical structure of the sentence.

The current study aimed to elucidate the importance of hierarchical structure during language processing using EEG. We recorded neural activity from 20 participants while they listened to streams of two-word sequences from four different conditions:

AN (adjective-noun): repetition of adjective-noun sequences,

AV (adjective-verb): repetition of adjective-verb sequences,

MP (mixed phrase): repetition of grammatical two-word phrases with varying grammatical categories,

RR (random): random word order; no phrases possible.

In the AN and AV conditions, grammatical categories occurred at a regular rate, so adjectives, nouns or verbs were repeated every other word. However the stream could only be parsed into grammatical phrases in the AN condition, for example:

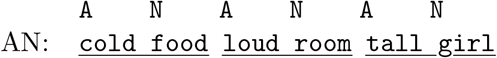

where underlining indicates grammatical phrases. In the AV condition, no such grammatical phrases could be formed, as in the example:

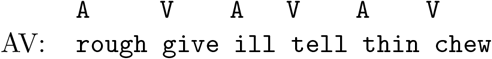

In the MP condition, grammatical two-word phrases could be formed, but grammatical category occurred with no regularity, for example

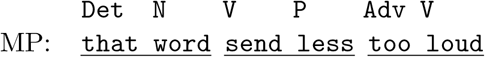

and, finally, in the RR condition there are neither grammatical phrases nor any regularity

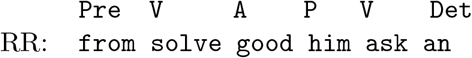

As in [7] the principle measure of response used here is the inter-trial phase coherence (ITPC); this quantifies the clustering of phases across trials at a given frequency. According to the hierarchical account we would expect a peak in ITPC at the phrase rate in both the AN and MP conditions, but no peak in ITPC in the AV or RR condition. A sequential account of language processing that relies primarily upon word-level statistics would instead predict a peak in ITPC at the phrasal rate in both the AN and AV conditions: our stimulus was designed so that the peak calculated using the fictive EEG simulated from word2vec gives similar phrase peaks in the AN and AV conditions.

Anticipating the main results, a peak in ITPC at the phrasal rate was significantly larger for AN than for any of the other conditions, suggesting that neural entrainment cannot be explained solely by hierarchical accounts or grammatical category regularity; rather, it additionally calls for higher level, syntactic representations. Our results support a language system that exploits both linear and hierarchical operations of language inputs to generate meaning.

## Results

The simulated EEG calculated from the word2vec representation yielded a peak in ITPC of the simulated EEG responses at the rate of syllable presentation (3.125 Hz, Fig. 2 in each of the four conditions. The model also yielded a peak in ITPC at the rate of phrase presentation (1.5625 Hz) for the AN and AV conditions (2), where, respectively, the grammatical adjective-noun phrases or the ungrammatical adjective-verb sequences were repeatedly presented. As described in the Methods, the stimuli for the AN and AV conditions were designed so that the vectors corresponding to successive words have similar distances in both of these conditions and the AN and AV conditions show similar peaks at the phrase frequency, even though one condition can be parsed into grammatical two-word phrases and the other cannot. The model also showed a pronounced, but lower amplitude peak in ITPC at the rate of phrase presentation during the MP condition. The phrase peak was absent from an version of the RR condition in which words are shuffled at random.

**Fig 2.**
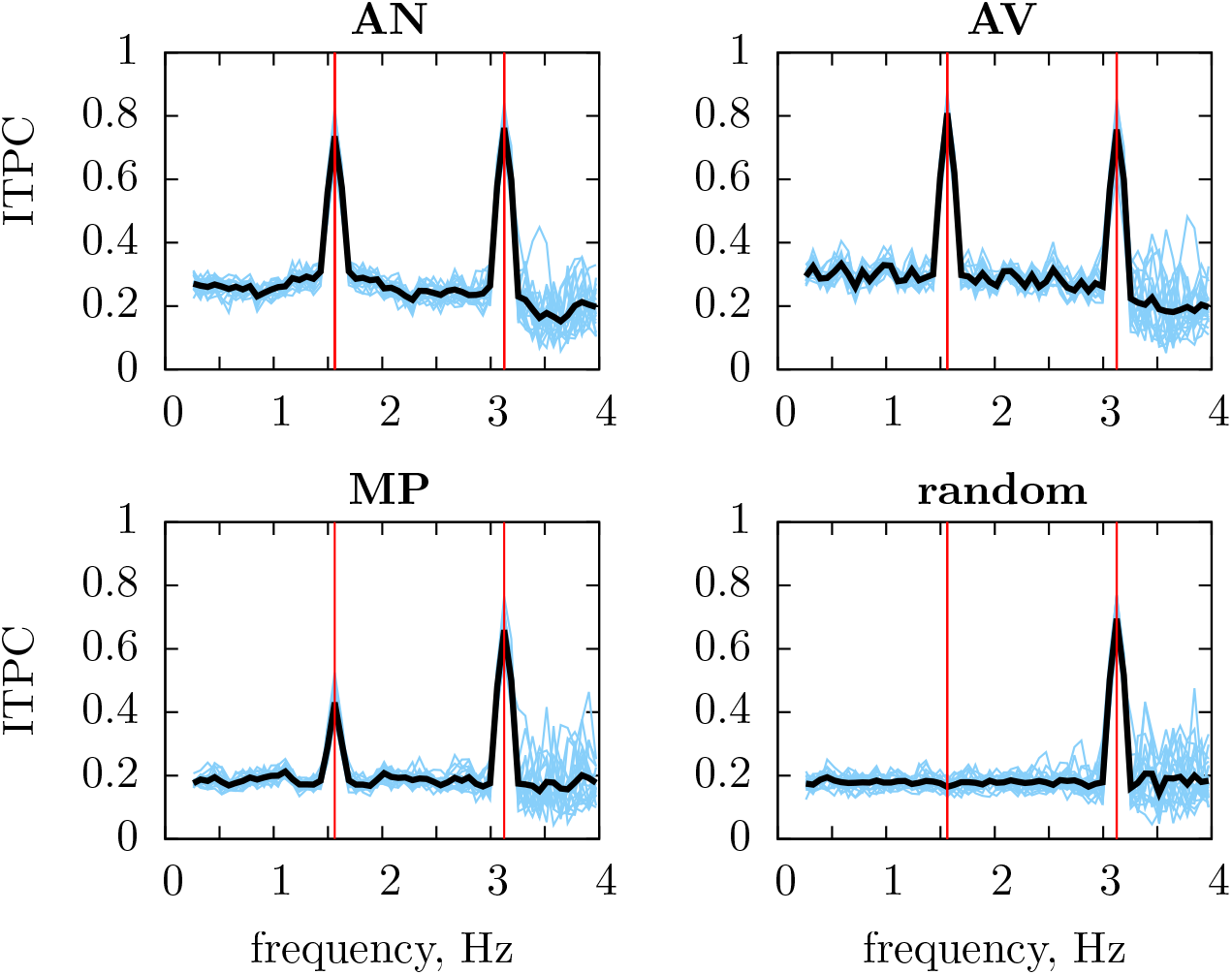
ITPC of the simulated EEG calculated from the word2vec representation for the four conditions. The model yielded peaks in ITPC at the rate of syllable presentation (3.125 Hz) in each of the four conditions (AN, AV, MP, random). Vertical red lines represent the frequency at which syllables and two-word sequences are presented, the blue lines show responses for 20 simulated particants, the black is the grand average over these participants.

Human EEG data showed a highly significant peak (*p <* 0.001) in ITPC at the rate of syllable presentation (3.125 Hz) in all conditions tested (Fig. 3). There was a highly prominent peak at the phrase rate (1.5625Hz) in the AN condition (*p <* 0.001), and a much less prominent yet still significant peak in the MP condition (*p* = 0.042). The phrase peak was not significant in the AV condition (*p* = 0.063). For the RR condition no evidence of a phrase-level response was found (*p* = 0.36). A reduction in the amplitude of ITPC peaks at the phrase rate in the MP condition when compared to the AN condition, and its absence in the RR condition, was consistent with the simulated data, but the less prominent peak in the AV condition was not.

**Fig 3.**
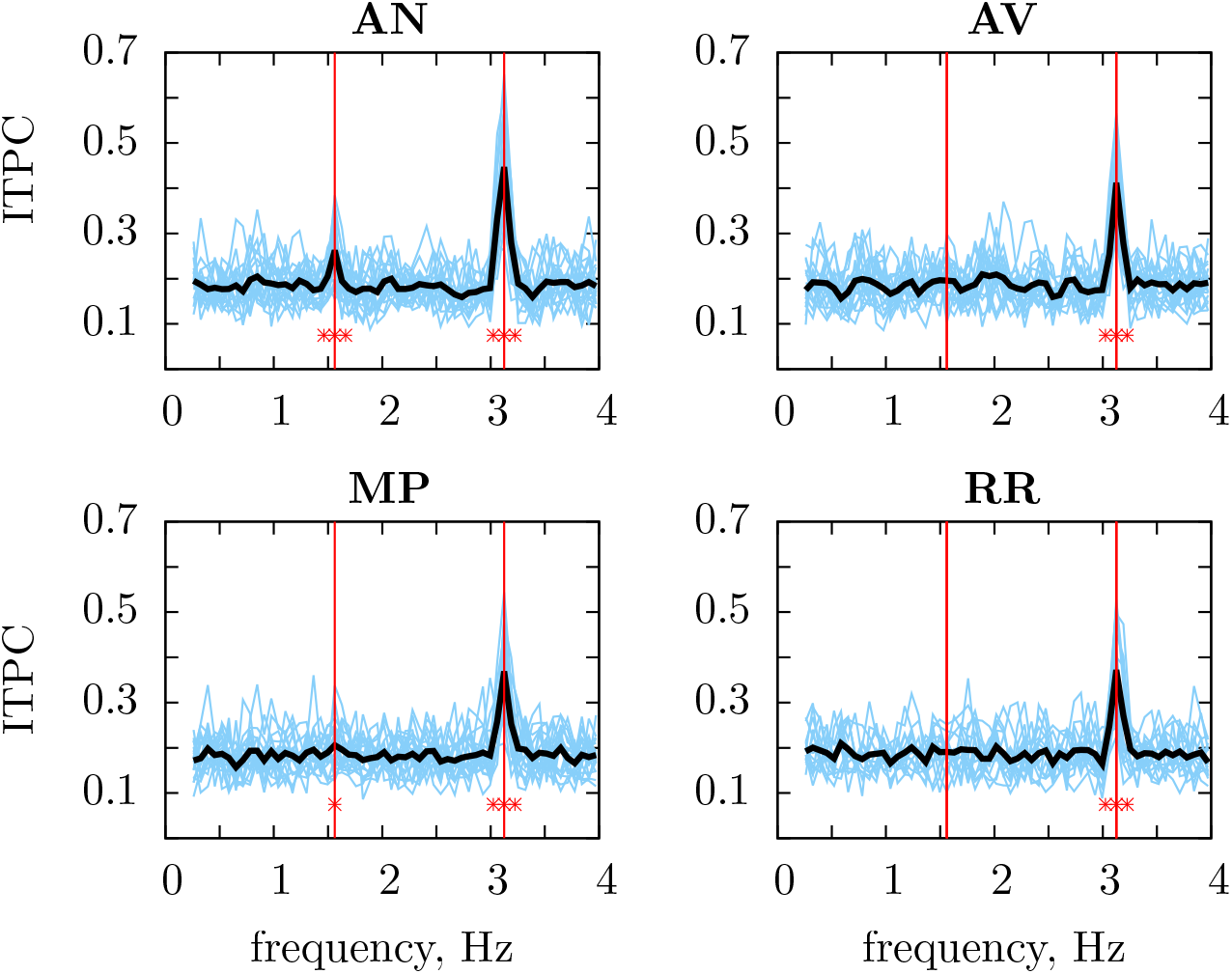
EEG responses recorded in human participants for each of the four conditions. Statistically significant peaks in ITPC values were observed at the rate of syllable presentation (3.125 Hz) in each of the four conditions (AN, AV, MP, RR). A statistically significant peak in ITPC is observed at the rate of phrase presentation (1.5625 Hz) in the AN and MP conditions. Red stars represent statistical significance *\** : *p <* 0.05, *\*\** : *p <* 0.01 and *\* * \** : *p <* 0.005. Vertical red lines represent the frequency at which syllables and two-word sequences are presented.

As described in the Methods section, data for the AN, AV and MP conditions come from 20 participants, there is only data for the control RR condition for 16 of these. Restricting the analysis to these 16 participants does not change the conconclusions from these results; on 16 participants the peaks at the rate of syllable presentatioon are significant (*p <* 0.001) for all participants; the phrase peak is significant for the AN condition (*p <* 0.001), the AV condition is significant (*p* = 0.044), the MP condition has *p* = 0.15 and RR, *p* = 0.38.

We have also performed an additional, ‘by item’ analysis with streams used as item, the ITCPs were averaged across participants to produce an average value for each stream for each condition. In this approach far smaller ITCP peaks were expected because of the likely differences in the phase of responses from participant to participant. In the ‘by item’ analysis the ITCP included both the variability of the response to stimulus, which we are interested in, as well as the less interesting variability in phase due to differences between the participants, for example, in their head shape and size. Nonetheless, the result was somewhat similar: there were significant peaks at the syllable rate (*p <* 0.001) and at the phrase rate for *AN* : *p <* 0.001; whereas the AV, AN and RR conditions showed no peaks.

In order to directly compare the ITPC values at the phrase rate across the four conditions, the Kruskal-Wallis test was used. The effect of condition was significant at the phrase frequency (*p* = 0.003). Pairwise comparisons (one-sided pairwise Wilcoxon signed-rank test, uncorrected) showed that the ITPC in the AN condition at the phrase rate was significantly higher than in the AV or MP condition (both *p* = 0.006); the difference (two-sided pairwise Wilcoxon signed-rank test) between the AN and MP condition was not significant (*p* = 0.82). The effect of condition on the frequency of syllable presentation had *p* = 0.051 and, perhaps surprisingly, the pair-wise two-sided tests indicate that AV was different from RR (*p* = 0.0042) and both AV (*p* = 0.041) and AN (*p* = 0.0025) were different from MP. This might indicate that the participants listened more attentively to the AN and AV stimuli.

Figure 4 shows individual particpants ITPCs for each condition. Statistically significant peaks at the rate of syllable presentation were observed in all participants in the AN and AV conditions, 19/20 participants in the MP condition and 15/16 in the RR condition.

**Fig 4.**
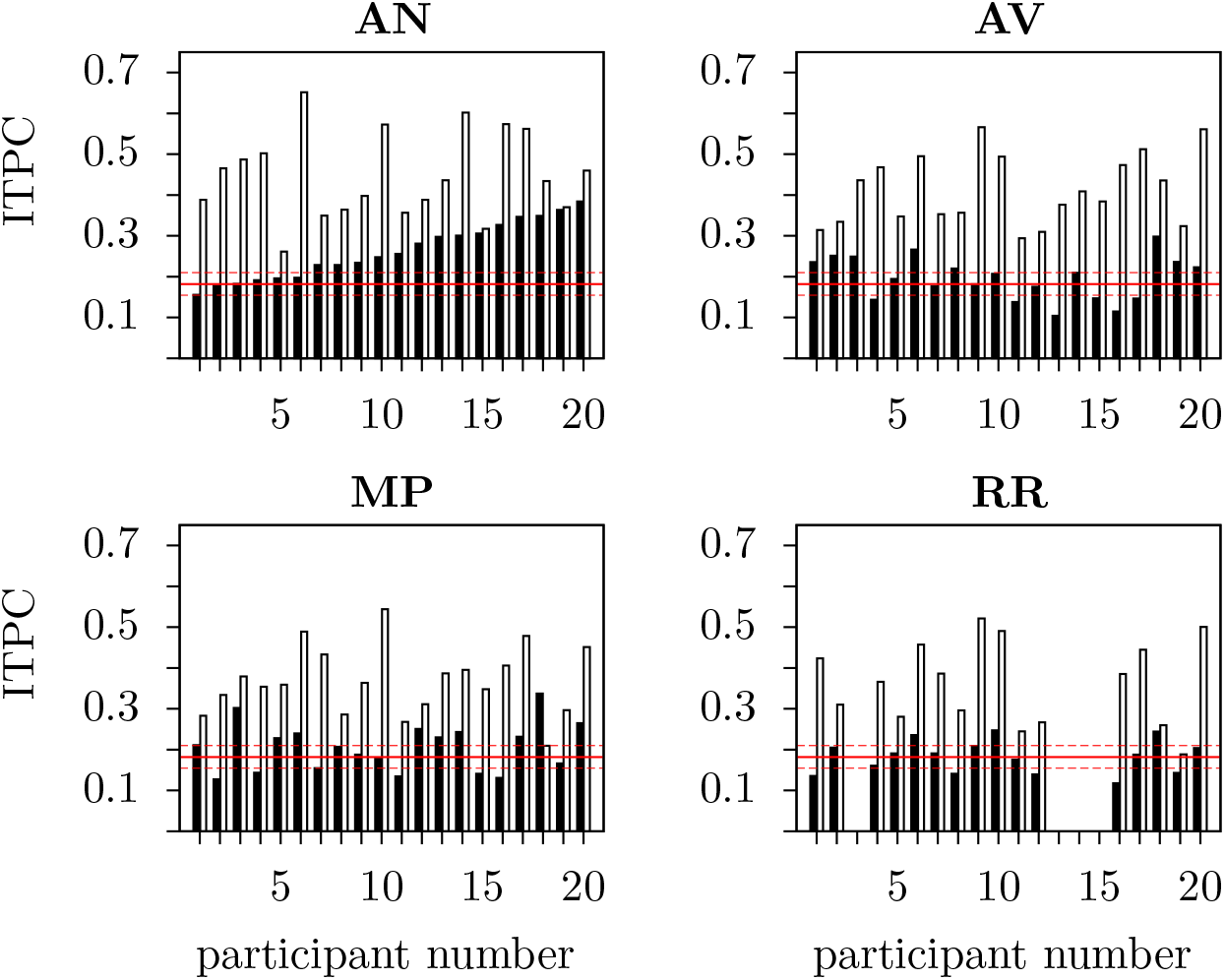
ITPC responses from individual participants at the phrase rate (3.125 Hz; filled bars) and the syllable rate (1.5625 Hz; unfilled bars) in each of the four conditions (AN, AV, MP, RR). For the AN, AV and MP conditions, values of the ITPC at each frequency of interest are displayed for each of the 20 participants, for RR, for the 16 participants. Participants one to 20 are ordered in accordance to their ITPC at the phrase frequency in the AN condition, in increasing order from left to right. Red horizontal lines indicate the mean ITPC value for random phases along with the corresponding significance thresholds (*p <* 0.05). These values are the same for both frequencies and so any ITPC that is above the upper red line is significantly higher than chance level.

When analysing the EEG responses of individual subjects at the rate of phrase presentation, a statistically significant peak was observed in 14/20 participants in the AN condition, in 7/20 participants in the AV condition, 9/20 in the MP and 3/16 in the RR conditions. Thus the pattern observed in the grand averages can be seen in the majority of individual participants.

## Discussion

The current study investigated whether, and to what extent, syntactic structure is automatically utilised by the brain during language comprehension, over and above information about grammatical category. It was found that repetitive presentation of grammatically well-formed, two-word adjective-noun phrases yields a prominent peak in ITPC at the rate of phrase presentation, but that repetitive presentation of two-word adjective-verb sequences that cannot be combined into a phrase did not produce a peak. The amplitude of the AN peak, as well as the syllable peak in response to single words, is consistent with previous findings [7]. This provides support for a syntactic operation that enables combining of words into higher-level syntactic units and suggests that the processing of linguistic input involves levels of abstraction beyond word-level grammatical-category information. This supports classical syntax-based approaches to language [2, 5, 9]. It is also generally compatible with the proposal that higher-level chunking of smaller language units occurs during language processing [6], although the nature of what the chunks are remains unclear. Indeed, in our analysis we have assumed that the grammatical categories we employed in designing our stimuli are relevant for language processing in the brain. There are other grammatical accounts that could be used to construct putative phrase conditions. Nonetheless, using a Chomskyan account of phrase structure has given us two conditions, AN and AV, which produced significantly different responses.

The distributional semantics model predicts a similar peak in ITPC at the rate of phrase presentation during both the AN and AV conditions. However, despite similarity in distributional vector space for the AN and AV conditions, the ITPC peak at the rate of phrase presentation was absent in the AV condition in the experimental recording. This suggests that the brain’s response is not merely a function of grammatical category; rather, it also reflects higher-level syntactic constituency.

The ITPC peak at the rate of phrase presentation was found in response to the MP condition was significantly smaller than the peak found in response to the AN condition, even though the MP condition contained repeated presentation of grammatically well-formed phrases. To the extent that each phrase involves combining two words into a single syntactic unit, there is a clear regularity in the MP condition at the syntactic level. This reduction might indicate that the response is not sufficiently abstract to reflect repetition of syntactic constituents independent of their lexical properties; for example, the phrases differ in the location of their head; determiner-noun phrases have the head after the modifier (that word) while verb-adverb phrases have their head before (send less). According to this interpretation, the phrase-level response found in the AN condition cannot be interpreted in its entirety as a reflection of the Merge operation [5] in its most general form: in both the AN and MP conditions words are pairwise merged into a phrase, but the ITPC peak is larger in the former case.

Another explanation for the reduction in phrase peak in the response to the MP condition is that streams for this condition are more difficult to follow and consequently less well attended to. To investigate this we performed a behavioural study in which subjects listened to a stimulus modelled on our EEG stimulus with streams for the AN, MP and RR condition. After the last stream they were asked to indicate whether they thought the last stream was composed of two-word phrases or random words. The order of the streams was randomized, thus each subject was only asked this question about one condition with a given subject had an equal chance of being asked about each condition: the AN, MP or RR stream; in fact there were 33 subjects asked about an AN stream, 27 about an MP stream and 28 about an RR stream.

For AN 32*/*33 *≈* 0.97 thought the stream was made of two-word phrases, for MP this was 24*/*27 *≈* 0.89 and for RR it was 22*/*28 *≈* 0.79. Thus, almost all subjects asked about an AN stream correctly identified that is was composed of phrases, but a substantial majority of subjects asked about an RR stream incorrectly believed it was in fact composed of phrases; MP lies between the two. A Fisher Exact test shows that AN subjects were significantly (*p* = 0.031) more likely to believe the stream was composed of phrases than RR subjects; the other two comparisons are not significant (AN*>*MP *p* = 0.234 and MP*>*RR *p* = 0.253). This behavioural experiment demonstrates the difficulty in parsing the stimulus as it is being listened to and does indicate that the MP streams are harder to distinguish from RR than the AN streams. See the Supplementary Information for a description of the design of this experiment.

A model of language comprehension is likely to exploit linear and hierarchical factors and describe how the brain uses different types of evidence: lexical, syntactic and semantic, in deducing meaning. While these different elements had seemed difficult to reconcile, recent neural network models with a linear temporal structure are able to discover and encode hierarchical structure, see [1, 14] for example. These models are consistent with the results outlined in the current study. Here we present evidence that words are combined by the brain into phrases and that syntactic information is important for the brain’s response to language; this indicates that hierarchical structure is deduced and classified, at least in part, based on syntactic information.

One issue with frequency tagging experiments, like the one presented here, is that neural mechanisms responsible for generating the ITPC in the frequency tagging paradigm are still not clear. It may be that cortical entrainment to the rate at which features of interest are presented causes the peaks in ITPC [15], or instead it is possible that the frequency tag is driven by regularities in ERPs in response to individual words or their combination. It is also possible that there is a variable ‘error’ signal associated with the irregularity of the MP condition and the ill-formed AV phrases and this is disrupting the response at 1.5625 Hz, the phrase frequency. Indeed, without a model relating neural dynamics to the ITPC we cannot be certain that the relative sizes of different responses are indicative of different manipulations of the incoming signal; this is a limitation of frequency-tagged experiments such as the one presented here.

In [20] it is argued that specific networks of neurons may be sensitive to sequences of words from discrete grammatical categories. In this way, local networks of neurons could learn to become sensitive to activation by a sequence of elements from different groups, for example an adjective followed by a noun, as in the AN condition presented in this study. This could explain why we see a larger ITPC at the phrase rate following repetitive presentation of phrases of the same type (AN). In other words adjective-noun sequences would repetitively and consistently activate the same sequence-detecting network of neurons responsible for processing adjective-noun sequences, whereas presentations of a mixture of different types of phrases would activate a different network of sequence-detecting neurons and thus generate an inconsistent EEG response and a smaller peak in ITPC at the phrase rate. On the other hand there would be no sequence-detector network for ungrammatical combinations, such as adjective-verb, and these therefore fail to elicit a phrase-level response.

In conclusion, the experiments described in the current study demonstrate that neural entrainment cannot be readily explained at the lexical level; rather, it additionally calls for higher-level syntactic representations. Yet, in our paradigm, frequency tagging of higher-level syntactic units emerged most strongly in the presence of grammatical category repetition, leaving open the question of how abstract syntactic representations are.

## Methods

### Participants

Twenty right-handed, native English speakers (12 female, mean age 25 years, range 22–42 years) participated in this study. Participants were screened for dyslexia and hearing impairments. All participants gave written, informed consent prior to undertaking the study and were reimbursed for their time at a rate of £10/hour. Ethical approval for our experimental procedures were obtained from the University of Bristol Faculty of Science ethics board. All methods were performed in accordance with the relevant guidelines and regulations.

### Stimuli

The experimental procedures were similar to those used in a recent EEG study [7]. Listeners were played streams of monosyllabic words in English. The words were synthesised individually using the `MacinTalk Synthesizer` (male voice `Alex, in Mac OS × 10.7.5`). All of the synthesised words (226– 365 ms) were adjusted to 320 ms duration and normalised in intensity using the freely available `Praat` software [3].

Monosyllabic words were selected from different grammatical categories, namely adjectives, nouns, verbs, pronouns, adverbs, determiners and prepositions. Words were only selected if they could be unambigouously categorised into a distinct grammatical category, so, for example words such as `drink, ride` or `walk` were avoided because they are ambiguous between verbs and nouns. All nouns were singular and all verbs were in the present tense.

The four experimental conditions were AN, AV, MP and RR:

1. Repetition of ‘adjective-noun’ sequences (AN).

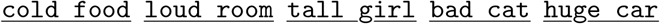 An adjective and a noun were repeated every other word. This condition contained grammatically correct two-word phrases, (underlined, with the grammatical category repeated every second word.
2. Repetition of ‘adjective-verb’ sequences (AV).

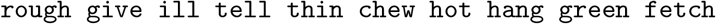 An adjective and a verb were repeated every other word. The word sequence in this condition preserved the repetition of grammatical category but did not contain grammatically well-formed phrases.
3. Repetition of grammatically well-formed phrases (underlined) without repetition of grammatical category information (MP).

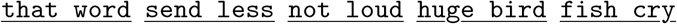 Grammatically well-formed, two-word phrases were composed from a pool of adjectives, nouns, verbs, pronouns, adverbs, determiners and prepositions. Phrases could take one of the following forms: ‘verb-noun’, ‘verb-adjective’, ‘adverb-adjective’, ‘determiner-noun’, ‘preposition-noun’, ‘verb-adverb’ and these were presented in a pseudo-randomised order to avoid repetition of grammatical category in adjacent phrases and to prevent grammatical phrases occurring across phrase boundaries: thus, for example, <u>not loud fish cry</u> would be excluded since <u>loud fish</u> is a noun phrase.
4. Pseudo-random word sequence chosen so that no phrases can be formed regularly between adjacent words.

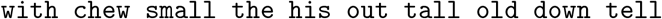 In this condition, words from the pool of adjectives, verbs, prepositions and determiners were randomly selected. Nouns were not included because they combine into grammatically correct phrases with words from many other grammatical categories.

A complete list of all stimuli used in the current study can be found in the Supplementary Information. Critically, in the AN and AV conditions, words were ordered such that there is not difference in similarity between the word2vec representation of consecutive words. Taking AN as an example, all the cosine similarities between the vectors representing adjective and nouns were calculated and only those pairs with values between 0.75 and one were retained; these values were hand-tuned to give a sufficient number of pairs while excluded as far as possible dissimilar pairs. To form a stream an initial pair was picked for this set, giving the first adjective-noun pair in the stream, *A*_1_ and *N*_1_. The adjective, *A*_2_ *≠ A*_1_, whose similarity to *N*_1_ is closest to the similarity of *A*_1_ and *N*_1_ is then picked. Next the noun *N*_2_ is picked so its similarity to *A*_2_ the closest to the similarity of *N*_1_ and *A*_2_. With the constraint that no pair appears twice, this is repeated until all 52 words are chosen. The same method was used to generate streams for the AV condition.

### Experimental Procedures

Each stream contained a sequence of 52 monosyllabic words played back to back in a continuous stream. Streams were therefore 16.64 seconds long. In total, participants listened to 150 streams, with 25 streams for each of the four conditions AN, AV, MP and RR, along with two filler conditions. An error in the marker file meant that one block was not usable, so 24 streams were included in the analysis. Blocks were made up of six streams and contained one stream from each condition plus the two filler streams. Within each block, streams were presented to the participants one after the other. After each stream, participants were asked whether they had heard any four word phrases, the instructions give three examples: <u>ask him this thing</u>, <u>from my old car</u> or <u>sit in that tree</u>. This acted as the attention trap with the four-word phrases occurring in ten percent of streams. These streams were not excluded from the analysis. Following the button press, the next stream was played after a delay of 250 ms. At the end of each block participants were given a 10s break, with a longer 2 minute break at the halfway point. The streams within each block were presented in a random order that was counterbalanced across participants but the composition of blocks and their order was the same across participants.

### EEG Recording

EEG signals were sampled at 1000 Hz from 32 Ag/AgCl electrodes fitted on a standard electrode layout elasticised cap using a BrainAmp DC amplifier (Brain Products GmbH). The EEG was recorded in DC mode, using a low-pass filter of 1000 Hz (fifth-order Butterworth filter with 30 dB/octave). FCz was used as a reference channel. The impedance of the electrodes was kept below 5 kOhms. Recordings were analysed offline using `MATLAB` (v. R2020b, Mathworks Inc.) and the `FieldTrip` toolbox (v. 20200607) [19]. As the recordings were performed using a 32-channel system (rather than a 128-channel system as, for example, [7]) we did not do dimensionality reduction on our EEG signals using PCA. Eyeblink artifacts were removed by applying ICA to the filtered signal. An independent component was removed if in its topography the mean power over the most frontal four channels (Fp1, Fp2, F7 and F8) was two times greater than the mean power over all other channels, as in [7]. As our signals of interest are in the low-frequency region, at 1.5625 Hz (phrases), and 3.125 Hz (syllables), the EEG signals were filtered offline using a 25 Hz low-pass filter (sixth-order Butterworth fileter with 36 dB/octave). Data were re-referenced offline to a common average reference. For each condition, individual streams (16.64s long) were epoched. Upon sound onset there is a transient EEG response and so the first four syllables (1.28 seconds) in each epoch were removed from the analysis. This meant that the overall length of the analysed part of each stream was 15.36 seconds (corresponding to 48 syllables × 0.32 s).

### Data analysis

After preprocessing, the EEG signal was converted into the frequency domain using the discrete Fourier transform with a frequency resolution of 0.0651 (1/15.36) Hz. The intertrial phase coherence (ITPC), what is also known as the square mean resultant, is

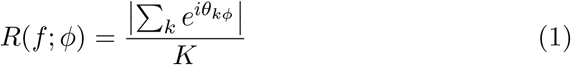

where *θ*^*c*^ is the phase angle of each complex-valued Fourier coefficient at frequency *f* and *k* is a trial index, with *ϕ* representing the other parameters such as the channel.

In most examples, the ITPC is calculated for each of the four different conditions for each participant and each channel; in this case *k* represents the different word streams corresponding to a given condition. In this case the ITPC is *R*(*f*; *pce*) where *p* labels participants, *c* conditions and *e* electrodes. This is averaged across electrodes to give *R*(*f*; *pc*) and, for example, the ITPC for different conditions is compared by examining the 20 pairs of values corresponding to the twenty partipants. For the ‘per-item’ analysis the ITPC is calculated for each condition for each stream and each channel so that *k* corresponds to the different participants. After also averaging across electrodes, this gives *R*(*f*; *sc*) where *s* is the index which labels the streams.

### Significance Testing

To determine whether a peak at one of the two target frequencies was significantly different from chance the ITPC was compared to the ITPC for random data. For the data an ITPC was calculated for each electrode using 24 phases computed for the 24 streams in each condition for the stimulus; this is then averaged over the 32 electrodes. To produce a simulated ITPC this calculation was mimicked for random phases. Thus, 24 phases were picked at random and used to calculate a ITPC for one ‘electrode’, this was repeated 32 times and the 32 values were averaged to give a simulated ITPC value which can be compared to the ITPC values calculated using the experimental data. To produce the confidence intervals for Fig. 4 5,000 of these simulated ITPC values were generated in this way, these were ordered and, for example, the 95% confidence interval corresponds to the 250th and 4750th entries in this list. To determine whether an ITPC peak was significantly different from chance a Mann-Whitney U-test was performed using these 5,000 values and the actual participant data: a Mann-Whitney rather then Wilcoxon test was used because these are not paired samples.

### Simulating Word Vector ITPCs

The word2vec repsententions for the words used in the stimuli were downloaded from (https://fasttext.cc/docs/en/pretrained-vectors.html). These were calculated using a distributional semantics model that was trained on a large English corpus [4]. Following [13] the simulated EEG was calculated from these vectors: the vectors are 300-dimensional so they give 300 channels. Time is discretized into 1ms quanta and a period of 320ms is allocated to each word. For a given stream let *v*_*e*_(*t*) denote the value of the voltage at time *t*. If **w**^1^ is the word2vec representation of the first word in the stream then for *t* ∈ [1, 320]

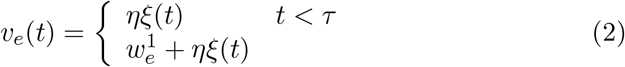

where *τ* is a delay chosen uniformly in the interval [20,60], *ξ*(*t*) is unit-variance zero-mean pink-noise and *η* = 0.5. This is repeated for each word in the stream, with independent *τ*. Individual participants correspond to a different random selection of 32 ‘electrodes’ from the 300 components and to different instances of the 1*/f* - noise: this is done to give the graphs some similarity to the graphs for the real data, but is not intended to model participant-to-participant variability.

The AN, AV and MP conditions use the identical stimuli as used in the experiment. However, the random condition differs from the RR condition in that the words are shuffled. In the RR condition adjacent words are chosen so as to not repeat grammatical category, the ITPC on these data is sensitive enough to detect this deviation from true randomness. With its different types of artificial noise, the simulated EEG is a complicated measure of the regularity of the stimuli. A much simpler measure is given by the Fourier coefficent

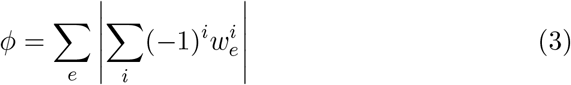

where 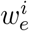 is the *e* component of the *i*th word in a stream. Averaging *ϕ* over streams and normalizing to the random condition gives values of 1.87, 2.41, 1.41, 1.13 for AN, AV, MP and RR respectively.

### Code and data availability

The data collected in this study is available at `doi:10.5281/zenodo.4019709;` the `Presentation 20.0` (Neurobehavioural Systems Inc.) script used to run the experiment, the stimuli and the code used for data analysis and for producing the simulated EEG is available at `doi:10.5281/zenodo.4275804.` In addition to the `MATLAB` code used to epoch the EEG data, perform blink-removal and calculate the Fourier transform, analysis and simulations were performed using `Julia` (v. 1.1.1). All data from the behaviour experiment and the scripts in `jsPsych` (v. 5.0.1) used to run the experiment are available online at `doi:10.5281/zenodo.4275815.`

## Acknowledgements

CJH and AB acknowledge support through a James S. McDonnell Foundation Scholar Award in Cognition (JSMF #220020239). NK acknowledges the support of the HSE RF Government grant 075-15-2019-1930. We are grateful to an anonymous referee for suggesting the behaviour experiment and Phillip Alday for suggestion the ‘per-item’ analysis and other improvements in our data analysis.

## Author Contributions

AB, NK and CH contributed to the conception and design of the work; AB carried out the experiments, AB and CH performed the analysis and prepared the figures. AB wrote the initial draft of the main manuscript text, CH and NK made a significant contribution during manuscript revisions.

## References

[1] M. Baroni. Linguistic generalization and compositionality in modern artificial neural networks. arXiv preprint 1904.00157 to appear in the Philosophical Transactions of the Royal Society B, 2019.

[2] R. C. Berwick, A. D. Friederici, N. Chomsky, and J. J. Bolhuis. Evolution, brain, and the nature of language. Trends in Cognitive Sciences, 17(2):89–98, 2013.

[3] P. Boersma and D. Weenink. Praat: doing phonetics by computer [computer program]. www.praat.org, 2019. Version 6.0.56, retrieved 20 June 2019.

[4] P. Bojanowski, E. Grave, A. Joulin, and T. Mikolov. Enriching word vectors with subword information. Transactions of the Association for Computational Linguistics, 5:135–146, 2017.

[5] N. Chomsky. The Minimalist Program. Current studies in linguistics series. MIT Press, 1995.

[6] M. H. Christiansen and N. Chater. The now-or-never bottleneck: A fundamental constraint on language. Behavioral and Brain Sciences, 39:e62, 2016.

[7] N. Ding, L. Melloni, A. Yang, Y. Wang, W. Zhang, and D. Poeppel. Characterizing neural entrainment to hierarchical linguistic units using electroencephalography (EEG). Frontiers in Human Neuroscience, 11:481, 2017.

[8] N. Ding, L. Melloni, H. Zhang, X. Tian, and D. Poeppel. Cortical tracking of hierarchical linguistic structures in connected speech. Nature Neuroscience, 19(1):158–164, 2016.

[9] M. B. Everaert, M. A. Huybregts, N. Chomsky, R. C. Berwick, and J. J. Bolhuis. Structures, not strings: linguistics as part of the cognitive sciences. Trends in Cognitive Sciences, 19(12):729–743, 2015.

[10] S. L. Frank and R. Bod. Insensitivity of the human sentence-processing system to hierarchical structure. Psychological Science, 22(6):829–834, 2011. PMID: 21586764.

[11] S. L. Frank, R. Bod, and M. H. Christiansen. How hierarchical is language use? Proceedings in Biological Sciences, 272(1747):4522–4531, 2012.

[12] S. L. Frank and M. H. Christiansen. Hierarchical and sequential processing of language. Language, Cognition and Neuroscience, 33(9):1213–1218, 2018.

[13] S. L. Frank and J. Yang. Lexical representation explains cortical entrainment during speech comprehension. PLOS ONE, 13(5):1–11, 2018.

[14] Y. Lakretz, G. Kruszewski, T. Desbordes, D. Hupkes, S. Dehaene, and M. Baroni. The emergence of number and syntax units in LSTM language models. In Proceedings of the 2019 Conference of the North American Chapter of the Association for Computational Linguistics: Human Language Technologies, NAACL-HLT 2019, Minneapolis, MN, USA, June 2-7, 2019, Volume 1 (Long and Short Papers), pages 11–20, 2019.

[15] L. Meyer. The neural oscillations of speech processing and language comprehension: state of the art and emerging mechanisms. European Journal of Neuroscience, 48(7):2609–2621, 2018.

[16] T. Mikolov, K. Chen, G. Corrado, and J. Dean. Efficient estimation of word representations in vector space. arXiv preprint 1301.3781, 2013.

[17] T. Mikolov, I. Sutskever, K. Chen, G. S. Corrado, and J. Dean. Distributed representations of words and phrases and their compositionality. In C. J. C. Burges, L. Bottou, M. Welling, Z. Ghahramani, and K. Q. Weinberger, editors, Advances in Neural Information Processing Systems 26, pages 3111–3119. Curran Associates, Inc., 2013.

[18] T. Mikolov, W.-t. Yih, and G. Zweig. Linguistic regularities in continuous space word representations. In Proceedings of the 2013 Conference of the North American Chapter of the Association for Computational Linguistics: Human Language Technologies, pages 746–751, 2013.

[19] R. Oostenveld, P. Fries, E. Maris, and J.-M. Schoffelen. Open source software for advanced analysis of MEG, EEG, and invasive electrophysiological data. Computational Intelligence and Neuroscience, 2011.

[20] F. Pulvermüller. A brain perspective on language mechanisms: from discrete neuronal ensembles to serial order. Progress in Neurobiology, 67(2):85–111, 2002.

