## Supplementary Information for "Grammatical category and the neural processing of phrases"

### Grammar, lexical category and the neural processing of phrases

### Supplementary Information

#### Words used to make the streams

##### Adjectives

- hot cold warm old bad good rich loud slow soft sad green sly shy full wise fine thin wide short ill rough calm loud low small tall huge large

##### Nouns

- rat tree dog man girl frog king car fact room word food bird skill wife thing cat sky lamp

##### Verbs

- send sing think want give tell ask weep sit chew hang solve fetch weep solve cry

##### Prepositions

- in at with from of up down on out

##### Determiners

- this that the him me an my his her

##### Numeral Nouns

- fish sheep

#### The stimuli streams

The streams marked in red were not used in the analysis.

##### Grammatical two-word phrases used in the AN condition

- old girl shy man wise thing loud room cold food large fact thin lamp tall sky soft cat bad word good dog rough skill huge tree small frog green bird sly wife rich rat full food ill king short car full sky sad king hot skill low lamp thin man sad rat

- large thing small frog green tree old dog good word bad cat soft food hot lamp wise girl rich fact rough sky huge man cold thing thin rat short bird sly wife shy skill low car sad room ill king full sky tall lamp loud fact large food cold dog loud room
- rich dog full sky thin bird hot car loud tree rough girl wise word soft thing cold man huge food good lamp small rat short dog tall room ill cat old king green wife bad frog sad skill rich fact low fact sly frog shy wife large girl shy man wise thing
- low sky rich man tall sky soft food hot lamp wise dog sad cat short rat thin tree loud skill large frog rough girl green car old room good wife sly bird cold fact low thing ill king huge word small frog shy wife bad lamp full king wise fact sad man
- rich lamp full sky thin bird hot car loud tree rough girl wise word soft thing cold man huge food good lamp small rat short dog tall room ill cat old king green wife bad frog sad skill rich fact low fact sly frog shy wife large girl shy man wise thing
- cold man rough dog shy cat sly skill huge room large food cold man thin thing soft word wise tree loud rat old bird rich fact ill girl small lamp low car sad king green wife bad frog full sky hot dog good word short girl tall frog loud sky green tree
- bad wife huge fact good skill sly thing shy bird loud dog cold food large room hot cat soft word wise tree rough girl rich wife sad car low lamp old king green man small sky ill rat full frog bad thing thin man short bird tall room loud dog cold food
- tall bird sly dog cold food large fact thin lamp tall girl good bird green tree old word bad cat soft thing rough frog huge car hot sky full skill small man low room wise rat ill king short wife loud thing sad girl rich fact shy bird loud dog cold food
- wise car thin fact large food cold dog loud room huge skill rough rat sad man ill wife old bird rich thing hot car green girl small lamp low sky bad tree short king good frog full cat wise word soft sky tall lamp sly tree shy room loud dog cold food
- cold bird hot dog loud room cold food large fact thin lamp tall girl good bird green tree old word bad cat soft thing rough frog huge car

sly king sad wife short sky small skill rich rat ill man wise thing shy bird low sky full word loud frog green tree

- cold wife small frog green tree old dog good word bad cat soft food hot lamp wise girl rich fact rough sky huge man cold thing thin rat short bird sly wife shy skill low car sad room ill king full sky tall lamp loud fact large food cold dog loud room
- rich cat low fact cold thing soft word wise skill loud tree thin rat short dog small bird large frog rough girl rich food hot lamp green sky tall room ill cat huge king good wife sad car sly man shy girl old man bad cat full bird loud dog cold food
- soft thing cold dog loud room large food small frog green tree old word good man ill fact rough cat short rat thin sky full skill tall car shy king rich lamp soft bird sad wife sly thing huge girl wise thing bad fact low skill hot king wise sky tall lamp
- rich cat low fact cold thing soft word wise skill loud tree thin rat short dog small bird large frog rough girl rich food hot lamp green sky tall room ill cat huge king good wife sad car sly man shy girl old man bad cat full bird loud dog cold food
- low wife soft sky tall lamp green bird good girl hot room huge skill rough rat loud fact ill food thin frog large dog short cat sad man cold thing rich wife sly car old king full word small tree wise thing shy bird bad skill low lamp thin man sad rat
- sly wife wise thing loud room cold food large fact thin lamp tall girl good bird green tree old dog bad word soft cat rough man huge sky hot car sly king sad wife short frog rich rat ill skill low fact small room shy tree full rat sly girl sad man
- small rat thin fact large food cold dog loud room huge skill rough rat sad man ill wife old bird rich thing hot car green girl small lamp low sky bad tree short king good frog full cat wise word soft sky tall lamp sly tree shy room loud dog cold food
- small fact low fact cold thing soft word wise skill loud tree thin rat short dog small bird large frog rough girl rich food hot lamp green sky tall room ill cat huge king good wife sad car sly man shy girl old man bad cat full bird loud dog cold food

- cold frog wise thing loud room cold food large fact thin lamp tall girl good bird green tree old dog bad word soft cat rough man huge sky hot car sly king sad wife short frog rich rat ill skill low fact small room shy tree full rat sly girl sad man
- low girl full sky thin bird hot car loud tree rough girl wise word soft thing cold man huge food good lamp small rat short dog tall room ill cat old king green wife bad frog sad skill rich fact low fact sly frog shy wife large girl shy man wise thing
- small sky ill wife old fact sly frog tall girl good bird green lamp hot food soft cat bad word wise skill loud tree thin rat short dog small room huge thing shy man rich sky low car sad king full sky rough girl cold lamp large cat shy dog rough skill
- hot rat good thing bad fact huge room large food cold dog loud bird shy cat sly skill rough rat sad man ill wife old tree small frog green sky wise lamp hot car low girl full king short word thin fact rich girl tall frog soft man old word good dog
- low room full sky thin bird hot car loud tree rough girl wise word soft thing cold man huge food good lamp small rat short dog tall room ill cat old king green wife bad frog sad skill rich fact low fact sly frog shy wife large girl shy man wise thing
- good cat tall tree green frog small food rich girl wise bird large skill shy rat thin sky full word short thing low room sly king sad car old lamp cold fact loud cat hot dog rough man huge wife bad thing good fact ill room soft rat sly girl sad man
- soft wife low fact cold thing soft word wise skill loud tree thin rat short dog small bird large frog rough girl rich food hot lamp green sky tall room ill cat huge king good wife sad car sly man shy girl old man bad cat full bird loud dog cold food
- green cat full sky thin bird hot car loud tree rough girl wise word soft thing cold man huge food good lamp small rat short dog tall room ill cat old king green wife bad frog sad skill rich fact low fact sly frog shy wife large girl shy man wise thing

**Ungrammatical two-word sequences used in the AV condition**

- ill fetch shy think loud hang sad ask sly chew thin sit cold weep rich tell huge fetch good solve wise give full want soft sing ill send large sit rough give bad hang hot want old fetch low weep green think tall ask short chew small tell wise solve good send

- low think small chew green hang sly send short fetch full sing bad weep rough want low think rich ask cold give old sit ill solve sad tell good think wise ask loud hang tall weep shy chew soft want thin tell hot sit huge sing large give good fetch huge solve

- bad sit rich tell huge fetch good solve wise give full want shy hang old weep cold think soft sing ill chew bad send tall ask short sit low chew large hang rough fetch loud send sad sit thin want hot weep green think small tell sly sing loud ask bad give

- small hang wise ask good give full want shy hang old weep cold think soft sing ill fetch rough tell small sit hot chew tall send bad solve huge think sad sing sly want loud tell short give rich hang large chew low weep green fetch thin solve wise send loud ask

- tall think old sing shy tell sly think cold weep rich fetch loud send wise sit sad ask bad give rough hang large chew low want ill solve short tell tall sit thin chew green hang small think full ask huge sing good send soft give hot solve wise fetch sly want

- low tell green chew thin sit cold weep rich tell huge fetch good solve wise give full want shy hang old sing sly think soft ask low send hot hang bad give rough sit large think short weep ill ask tall fetch small want sad tell loud chew loud send wise sing

- shy ask sad weep loud sing sly want huge think shy tell tall sit large chew low hang short give rich fetch good solve wise send ill ask full give bad hang hot want thin tell old fetch rough think soft sing small send cold ask green solve wise weep sly chew

- full chew sad weep loud sing sly want huge think shy tell tall sit large chew low hang short give rich fetch good solve wise send ill ask full give bad hang hot want thin tell old fetch rough think soft sing small send cold ask green solve wise weep sly chew

- small sing hot hang old weep cold think soft want full sit sad send ill tell thin chew sly ask shy sing wise fetch good solve huge give short tell tall sit large think rough give bad hang rich want low weep green fetch small send loud ask wise solve good sing

- bad chew ill weep good fetch huge tell rich give short think rough sit large chew low hang wise send loud ask bad want sad sing sly solve hot hang old weep cold think soft want full sit thin chew green give tall ask small solve shy send wise sing soft fetch

- old solve low chew wise hang short sit huge sing ill fetch shy give sad solve large weep rough want cold tell small think full ask sly send rich tell tall sit thin chew green hang soft think hot weep old sing good send loud fetch bad solve wise give full want

- rough solve green chew thin sit cold weep rich tell huge fetch good solve wise give full want shy hang old sing sly think soft ask low send hot hang bad give rough sit large think short weep ill ask tall fetch small want sad tell loud chew loud send wise sing

- tall solve rough give bad hang good solve huge tell rich weep cold think soft want full sit sad send ill sing tall chew hot fetch large ask green chew thin sit wise hang short weep old tell small sing low ask sly fetch loud send shy solve wise give full want

- tall want green chew thin sit cold weep rich tell huge fetch good solve wise give full want shy hang old sing sly think soft ask low send hot hang bad give rough sit large think short weep ill ask tall fetch small want sad tell loud chew loud send wise sing

- green ask large sit rough give bad hang good solve huge tell rich weep cold think soft want full ask ill send sad chew old fetch loud sing sly weep wise give short think hot tell thin want shy hang low chew green sit small sing tall fetch wise solve good send

- thin send rough give bad hang good solve huge tell rich weep cold think soft want full sit sad send ill sing tall chew hot fetch large ask green chew thin sit wise hang short weep old tell small sing low ask sly fetch loud send shy solve wise give full want

- cold weep rich tell huge fetch good solve wise give full want shy hang old weep cold think soft sing ill chew bad send tall ask short sit low chew large hang rough fetch loud send sad sit thin want hot weep green think small tell sly sing loud ask bad give

- cold sing green chew thin sit cold weep rich tell huge fetch good solve wise give full want shy hang old sing sly think soft ask low send hot hang bad give rough sit large think short weep ill ask tall fetch small want sad tell loud chew loud send wise sing
- green send low chew wise hang short sit huge sing ill fetch shy give sad solve large weep rough want cold tell small think full ask sly send rich tell tall sit thin chew green hang soft think hot weep old sing good send loud fetch bad solve wise give full want
- soft solve loud weep sad think huge want sly sing shy tell tall sit large chew low hang short give rich fetch good solve wise send ill ask full give bad hang hot want thin tell old fetch rough think soft sing small send cold ask green solve wise weep sly chew
- full give loud weep sad think huge want sly sing shy tell tall sit large chew low hang short give rich fetch good solve wise send ill ask full give bad hang hot want thin tell old fetch rough think soft sing small send cold ask green solve wise weep sly chew
- low sit ill weep good fetch huge tell rich give short think rough sit large chew low hang wise send loud ask bad want sad sing sly solve hot hang old weep cold think soft want full sit thin chew green give tall ask small solve shy send wise sing soft fetch
- Probe trials
- loud solve soft chew shy weep tall sit large think hot tell ill ask full send old want cold solve sad give low fetch thin sing rough hang short tell huge fetch good solve wise give loud send sly hang green sit rich ask small weep bad chew loud want wise think
- small chew green chew thin sit cold weep rich tell huge fetch good solve wise give full want shy hang old sing sly think soft ask low send hot hang bad give rough sit large think short weep ill ask tall fetch small want sad tell loud chew loud send wise sing
- shy hang old sing shy tell sly think cold weep rich fetch loud send wise sit sad ask bad give rough hang large chew low want ill solve short tell tall sit thin chew green hang small think full ask huge sing good send soft give hot solve wise fetch sly want

**Grammatical two-word phrases used in the MP condition**

- too good want fame from rice sit more want milk sheep give solve less her cat not loud not thin slow thing sheep jump cold rat from fame the dog chew more her room sheep give chew hair in fame my sky fish want give well give sand soft rat of fame

- soon rich want wine this word fish cry my car in sand shy bird send sand sing soon fish weep want sand send less chew rice too short from salt fish drink fish make rich man warm tree from art too old less bad loud man that man ask soon from hair

- fish cry chew rice fetch less too cold on milk this fact my wife sheep think this bird want salt fish think green tree of wine less low this wife chew sand too small in wine wide tree ask well loud sky soft thing sing fast sheep cry sing far send art

- sheep laugh send not her skill fish laugh fetch fame want hair fetch art on art want milk his fact too old in sand fetch soon rough cat this girl large girl ask soon fish go loud cat sing more sheep think less calm of milk that girl from salt more loud

- chew hair think more less old sheep solve with sand with wine warm tree send not bad girl chew hair send far sheep give want gold her car sit too not soft in wine hang rice the sky his word fish make more warm this tree less green slow frog in wine

- of fame warm dog my lamp want rice give fast with rice give well not low fish solve soon cold want sand tell more want art fish think fish go with sleep less slow give salt this girl on gold loud word loud wife calm word that lamp her tree less tall

- want sand from gold of gold that skill sheep want in sleep fish give short dog too cold fetch soon this skill want milk that lamp too slow tall girl fish go fetch fame his cat cold room fetch less less huge rich wife fish weep on sleep think less fetch rice

- ill wife slow girl more calm give less want gold of rice not loud fish want the food ask fast fish sing fish eat give fast wide wife solve less his fact not tall too rich small king my food send sand give sand in fame sheep make want salt with rice

- fine food with hair weep more slow wife that lamp fish jump too hot send far give more want fame fish make fetch salt too calm in milk wide dog send art on wine fish sing want art too calm the girl fine room sheep give from wine my thing too slow
- sheep weep more rich want salt the lamp fish weep soon shy fine room rich dog hang art full wife fish cry not low from gold his room chew more fetch fast want salt want more from salt of gold her word that lamp of sleep send milk sheep solve sit soon
- send wine send art the wife not fine rich thing fetch gold sing well on sleep my girl her word from sand with milk sheep cry wide thing sheep cry soon cold too hot sheep sing not slow sing well fish weep her car want sand chew fast huge dog in fame
- less fine not large with sleep loud sky want sleep chew wine solve more want wine cold rat my food my king send fast solve fast soon rough on wine fish go her wife fish jump sit fast in salt fetch rice with sleep sheep laugh the food sheep laugh not ill
- think well green room fish jump his girl good thing more warm too loud tell less fetch art of gold sheep weep of art this skill send wine sheep solve want art with rice sly wife sheep go more low this girl from sand her wife want more thin cat less large
- send fame fish want give salt green dog the lamp give wine wide lamp in art the man sheep go not slow of sleep on salt give milk fish make fish solve fine skill in rice give well her bird weep more chew fast her rat too cold hang far slow fact
- on rice from rice more fine give sand ask soon low sky hang fast sheep eat the word send salt solve less sheep sing on rice his sky too short too cold sly man on rice the dog not old send fame my man huge man want rice fish cry weep not
- fish cry this sky too rough wise thing hang rice want sleep with gold on art give wine want salt bad man hang less shy cat sit more want more sheep solve this fact not thin from gold that fact that sky sit well fish make less wide sheep give fine fact
- cold girl sheep want in sand sit less want rice give fast too large less hot send gold sit fast good wife his car her wife fish drink from wine

his fact fetch rice sheep give huge man not calm from fame soon thin on wine soft room chew wine send fast

- sad man send wine on art slow food fish think of sleep hang art less old my sky send wine on wine want not fish weep not shy tell less this tree send wine sheep sing sheep give not bad soft frog this fact fine lamp my dog of fame sit too
- fish give this skill this room not shy fetch sleep large frog give soon with milk in sand less cold think more with hair fish go his skill the skill chew less fetch more rich man small car send rice old skill sheep go fetch fame from art less loud fish make
- fish laugh short word not ill wide sky the dog hot rat soon fine give soon this car low lamp that bird fetch milk too large give wine sit fast from salt send less hang rice of milk want gold her skill of wine soon wide in sand fish sing fish drink
- not sad sheep give sit too hot king want salt not short fish cry that word want art with gold from gold bad cat in sleep too loud that word soft king on sleep sheep want too bad fetch fame send less that food sheep go huge bird want hair sit soon
- not full more green that king hang fast weep more the room fetch wine calm wife her word in rice send more rich man too short sheep want with hair sheep laugh that rat sing far of rice in sleep sad wife fetch sleep warm tree hang rice sheep sing fetch sand
- Probe trials
- this car of hair not loud give rice more wise not shy warm room calm cat fish solve from salt of hair sheep give her rat fish make solve fast ask soon too rich in art sheep drink tall cat chew rice chew wine his wife fetch art good skill my word
- fish make this dog her frog the dog on art less short want fame fish cry sheep give cold room less loud less low of gold sing less fish cry chew rice her cat soon full good girl give sand of salt send soon with hair want rice green thing sit not
- of sand fish laugh her word with rice wide sky this thing chew sand fish jump think well that car more low less full sheep weep sheep solve from fame more loud more sly hang art from sand that sky huge room fetch soon tall lamp small food give salt fetch gold

**Ungrammatical two-word sequences used in the RR condition**

- tall in fetch her shy him down want my good at think from weep ill the solve this on sly sad me send down sing of old that hang him green of give an with short from tell full his hot my up sit out ask low the large her chew with

- up want ill an sing her in old this small out weep low his chew from cold me down fetch ask with hot him sad on send him solve the soft out give in sly that loud the from tell his green at think shy my sit of hang an rough down

- from solve good him ask an of rich his cold down want low this sing on huge my with sit hang in rough me full down think the give that small in weep out thin my soft me at chew her ill up send hot him tell on fetch that shy out

- of sing wise her solve an in huge this small with fetch full my send in sly the on want give up hot him loud at think that chew him old from tell up green this bad me at ask his rich down sit thin an hang down weep the short out

- old on send that rough an with ask the low from solve up chew rich me hang an down sad large my want with sing at bad him think me wise up fetch that out loud in sit shy his cold this of tell on give sly his thin her weep down

- me soft on sit old out sing the chew that down green hang him rich with wise this down solve in send thin his fetch from small an full her give at his hot on ask with tell rough the think that sad up shy of weep him want of tall my

- loud at tell an tall that in send her thin up ask down weep low my fetch her of wise short an think from give on huge his want that old down sing him in green with sit full the small the out hang at chew ill this soft me solve up

- want the wise down loud that at weep huge up tell her think up large him ask his at shy from chew rough me low an give with her green of fetch old me in sing on hang small my solve in tall the cold my sit out this thin down send

- with chew small the sing his out tall that old down tell soft him think at loud an of weep ask at low the thin from sit my fetch me green down hang on bad her sad this in solve my rich up give wise me want from send an good with

- good with hang my hot that up sit this old at think of sing large his chew her on rich small an tell in send of bad him weep me tall out solve her at thin down give loud an shy him with want from fetch ill that rough the ask from

- my sly down weep small of sing him give her from shy send that good in thin an down fetch at sit soft this tell up low me rough the think with him ill out solve on hang old his chew her sad from large of want that ask with huge this

- at weep bad that ask an down thin him cold of give huge that sing on loud her down sit hang with good me soft out fetch this solve this shy of think with rich her sad my up send his short out tell green the want in chew the sly from

- up send old me chew that at tall my rough out weep loud his sing down shy the of solve think at green him wise in tell the fetch an cold in hang with sad this bad her down want his small from sit thin my give with ask this rich on

- large out fetch her cold this in tell me short up give of think small me sit an with bad sly an hang up ask at wise the chew my rich at send him down hot with want low that shy his on solve of weep green the thin him sing from

- weep this short from rough my down chew full on ask his fetch of soft an give her in low out solve small me wise an hang down that bad up sit rich this out send with want large the sing at sly that old him think up me huge on tell

- this full up hang wise out send my give me in short sit an shy at old this of ask down fetch ill the tell from thin an hot her sing from my low on chew up solve green that weep his tall at loud on want him think with large the

- soft up sit him bad an with fetch this large in weep down hang huge the solve this on small green the chew from send with short her think him loud in tell that at low from give ill her old me out ask of want wise his sly my sing out

- out sing rough her want my from soft her cold down sit good me fetch up full him on send hang of green the bad up chew him ask that sad down tell at short his thin an of solve the ill in think small this weep with give that huge out
- wise of fetch an old that in ask him huge out give on tell cold my want the at large low the sit from chew from sly his send her soft with solve me on rich with sing short this tall an up hang down weep thin her sad that think at
- this full with ask soft down hang that chew an of rich sing the sly on tall her on give out send old this weep in bad that cold an want with his wise out fetch down think low him solve her loud up sad from sit my tell at small me
- weep her old at large me from want thin with sit his send on sad this sing me down sly of tell good an hot that solve at his wise in think short him out fetch from ask full the chew out soft the bad that give up my cold in hang
- huge with tell his tall the in sit her large out weep at give sly this ask him at low good him sing up hang on bad that think an shy from chew my with wise down want short his green an on fetch out send cold the thin me solve of
- Probe trials
- want my loud down old an out weep ill up hang that think of low him solve her from sad down tell hot him sly the ask at that good with chew huge her in give in fetch tall his send on soft this rough my sit with me large on sing
- rich down solve an cold that in hang me rough down fetch from ask sad the sing this out ill good that send from sit at loud my think her full in tell him of thin out give hot her soft his with want on chew shy him green the weep up
- ask an small in hot this out fetch wise out send the hang in thin the solve me of shy on chew short him sly my want with an old with sing full him down tell from sit rough his give of bad that huge her weep at this low up think

##### Instructions used in the EEG experiment

You will listen to 150 word streams.

The streams of words don't always make a lot of sense. Some streams will make more sense than others.

At THE END of each stream of words you will be asked whether you heard any phrases of at least four words.

Three example phrases are:

'ask him this thing'
'from my old car'
'sit in that tree'

If you think you heard a four word phrase please press 'y'
at THE END of the word stream and ONLY WHEN PROMPTED.

If you didn't hear any four word phrases please press 'n'
at THE END of the word stream and ONLY WHEN PROMPTED.

#### Behavioural experiment - methods

For the behavioural experiment four streams were chosen for each of the AN, MP and RR conditions along with one probe stream for each of the same conditions. The probe trials all contain a four-word phrase. For each subject the streams were shuffles and two probe streams picked randomly from the four where inserted into the list at locations five and 11, making fourteen streams in all. During the experiment the streams were played in sequence and after each stream the subject was asked "Did you hear any four-word phrases?". After the final stream they were also asked "In the last stream of words you hear any two-word phrases?" and "Write down as many things from the last stream of words you have just heard as you can remember".

90 subjects were recruited on prolific academic (`prolific.ac`) with a 1GBP payment. The subjects were pre-screened by prolific academic as United Kingdom or Irish citizens living in the United Kingdom or Ireland and were asked to identify their first language; two subjects who did not answer English were excluded. No other demographic information was collected.

The average number of words remembered from the previous stream was very low; on average subjects recalled just 3.82 words on average and often these words were not from the most recent stream or even not represented in the experiment at all. There was also no significant difference between the three conditions in the number of subjects who mistakenly answered that the previous stream contained a four-word phrase.

#### Extra Figures

A B

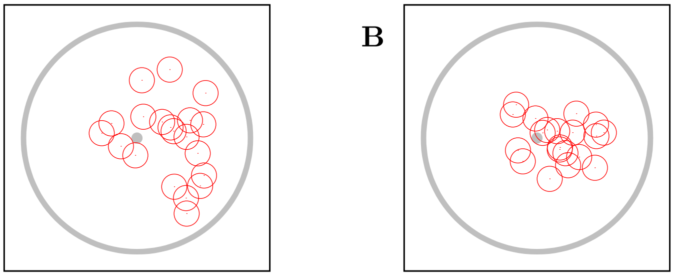

Figure 1: **The mean phases for the AN condition recorded at electrode TP9**. As an example of how the phase varies from participant to participant the mean phase over trials is calculated for each participant for electrode TP9, the electrode with the largest average ITPC at the phrase rate. The ITPC is calculated using the absolute value of $z_{pf} = \sum_s e^{i\theta_{kpf}}/S$ where $s$ indexes the different word streams and $S = 24$ is the number of streams for each condition, $p$ is the participant index and $f$ is the frequency. In **A** $z_{pf}$, red circles, are plotted for the response at the syllable rate: $f = 3.125$ Hz; the grey dot marks $z = 0$ and the grey circle $|z| = 1$. **B** is the same plot except it shows the response at the phrase rate: $f = 1.5625$ Hz.

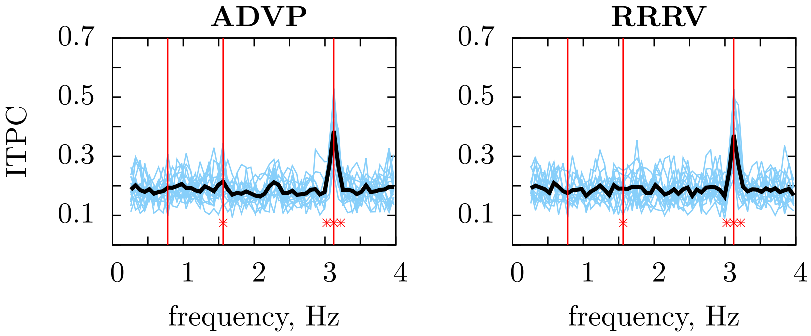


Figure 2: **ITPC for the extra conditions calculated using 16 participants**. The experiment had two conditions not included in the analysis; these conditions are decribed as fillers in the Methods section. These conditions were designed to investigate the 'sentence level' response. These streams have 12 four-word phrases instead of 24 two-word phrases, the phrases are ungrammatical but with the repetition of lexical category. In the RRRV condition the phrases includes three random words which are not verbs followed by one that is; in the ADVP condition the phase has the form adjective-determiner-verb-pronoun. It was hypothesised that because of the repetition of lexical category there would be a peak at 0.78125 Hz. In fact, there is no evidence of this peak. There is a peak at the syllable rate ($p < 0.001$) and peaks (ADVP: $p = 0.027$, RRRV: $p = 0.018$) at the phrase rate.
